# Mashona Mole-Rat Automatic Individual Identification Based on the Mating Calls

**DOI:** 10.1101/129452

**Authors:** Veronika Dvorakova, Ladislav Ptacek, Ema Hrouzkova, Ludek Muller, Radim Sumbera

## Abstract

In this study was tested mole-rat vocalization for presence of diverse individually distinctive features. An automatic system based on the GMM-UBM was used for individual recognition. The system distinguishes the recordings of the five mole-rats females. The overall achieved identification accuracy is 76.7%, the lowest 59.2%, and the highest 83.5%. The overall percentage is thus high enough to prove that the mating calls of the Mashona mole-rat can carry information about mole-rat individuality. Our results showed that studied vocal signals in the Mashona mole-rats are individually specific which indicates the possibility of individual vocal recognition in this species.

## Mashona Mole-Rat Automatic Individual Identification Based on the Mating Calls

### Introduction

The zoologist of the University of South Bohemia, Faculty of Science keep large number of underground rodents in a breeding facility. Because the mole-rats live in the soil under the surface, its vocalization is very important for communication. Many experiments were performed to reveal the detail of its vocalization, communication, and the capability of the mutual recognition. For example naked mole-rats (*Heterocephaus glaber*) use vocalization to distinguish size and dominance status of an individual, to stimulate a mate sexually, or to warn against danger (Yosida et al. 2007; Yosida and Okanoya 2009). Purpose of our experiment was to prove a possibility of the mole-rat individual identification based solely on its mating calls.

### Methods

The GMM-UBM method (Reynolds 2000) is adopted from the well-known Speaker Recognition task, which is ordinarily used in human speech, as well as in animal recognition research. The introduced system is tailored for individual identification on the open set, even when using non pre-processed recordings. The system performs identification with a sequence of particular verification trials.

A new framework tool was created in Matlab (Mathworks Inc. 2010), based on the GMM-UBM. This framework is designed for bird individual identification task (Ptacek 2016), and was re-designed for mole-rats identification. Figure 1 displays an outline of the GMM-UBM recognition system.

**Figure 1:**
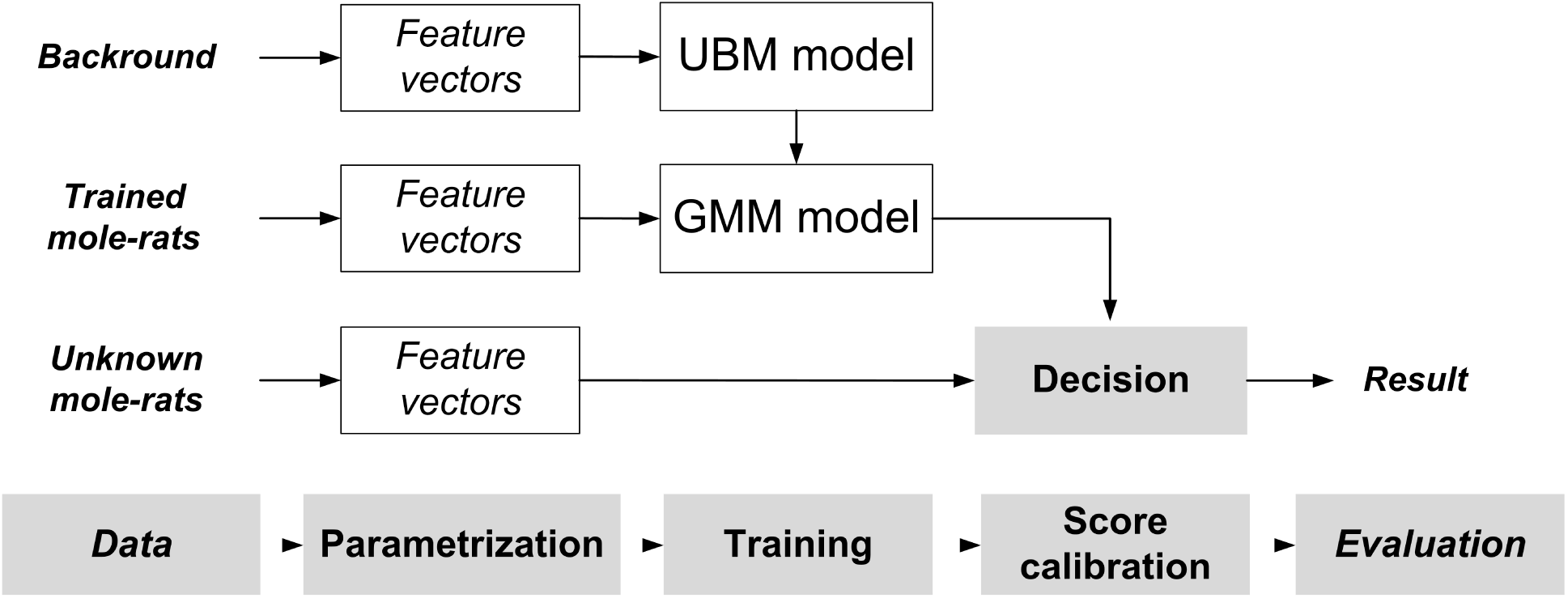
General outline of the GMM-UBM recognition system.

The process flow is decomposed into:

- Parametrization, feature extraction: recordings are parametrized to extract features, forming a set of feature vectors.
- Training: UBM model and GMM model estimation.
- Decision: probability comparison of unknown bird and trained models.
- Score calibration: choice of a verification threshold.
- Evaluation: based on EER, DET or any other method.

Mole-rat vocalizations are parametrized to extract relevant features, forming a set of feature vectors called Linear Frequency Cepstral Coefficients (LFCCs). In order to extract multiple feature vectors, a sliding window is utilized. After samples in the window are processed, the window is shifted to the next position, usually by half of its length, and the extraction of feature vectors is repeated. Samples in the sliding window are first weighted by a Hamming window to suppress undesirable effects, see Figure 2. Then, the Fast Fourier Transform (FFT) is applied right after the windowing. Next, the power spectrum is computed in order to extract frequency characteristics of the signal present in the window. To smooth the spectrum, a set of triangular shaped Filter Banks (FBs) – band pass frequency filters – of height one is spread across the frequency domain. After the filtering, the cepstral coefficients are calculated. Finally, in order to incorporate some dynamic information on the variation of the signal in time, delta coefficients are evaluated and added at the end of the cepstral vector (Bimbot 2004). Once the feature vectors in the form of LFCCs is extracted, the next step consists in modelling of the probability distribution of the data. The GMMs was firstly introduced to the speaker recognition by (Reynolds 1995) and is widely used up to now. GMMs are generative statistical models, well suited for description of static (context-independent) data sources, where the time progress of samples is of no interest. An example of a GMM is depicted in Figure 3.

**Figure 2:**
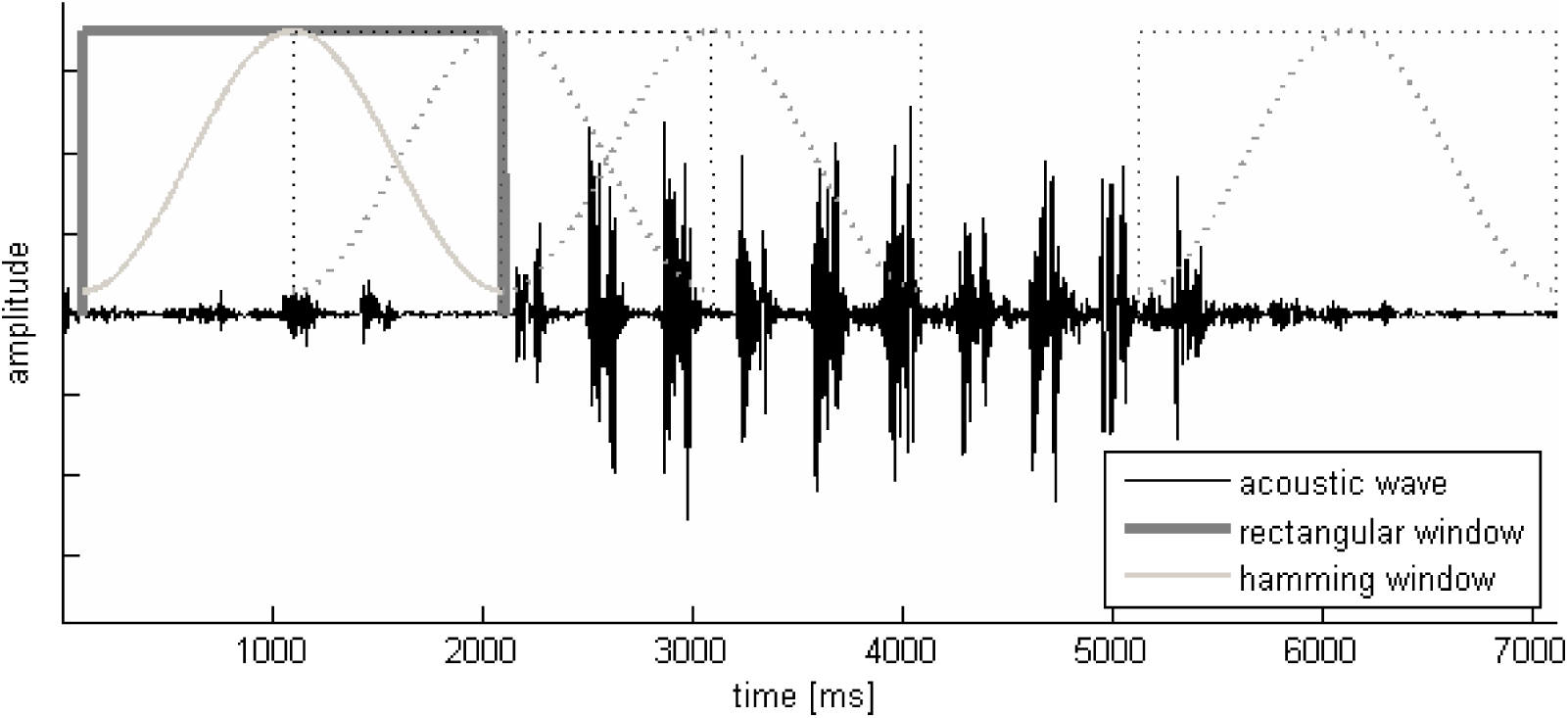
Samples in the rectangular window are weighted by the Hamming window, FFT is performed, filtration utilizing triangular filters is carried out, and a cepstral feature vector is extracted. Subsequently, the window is shifted to its new location and the extraction process is repeated (Machlica 2012).

**Figure 3:**
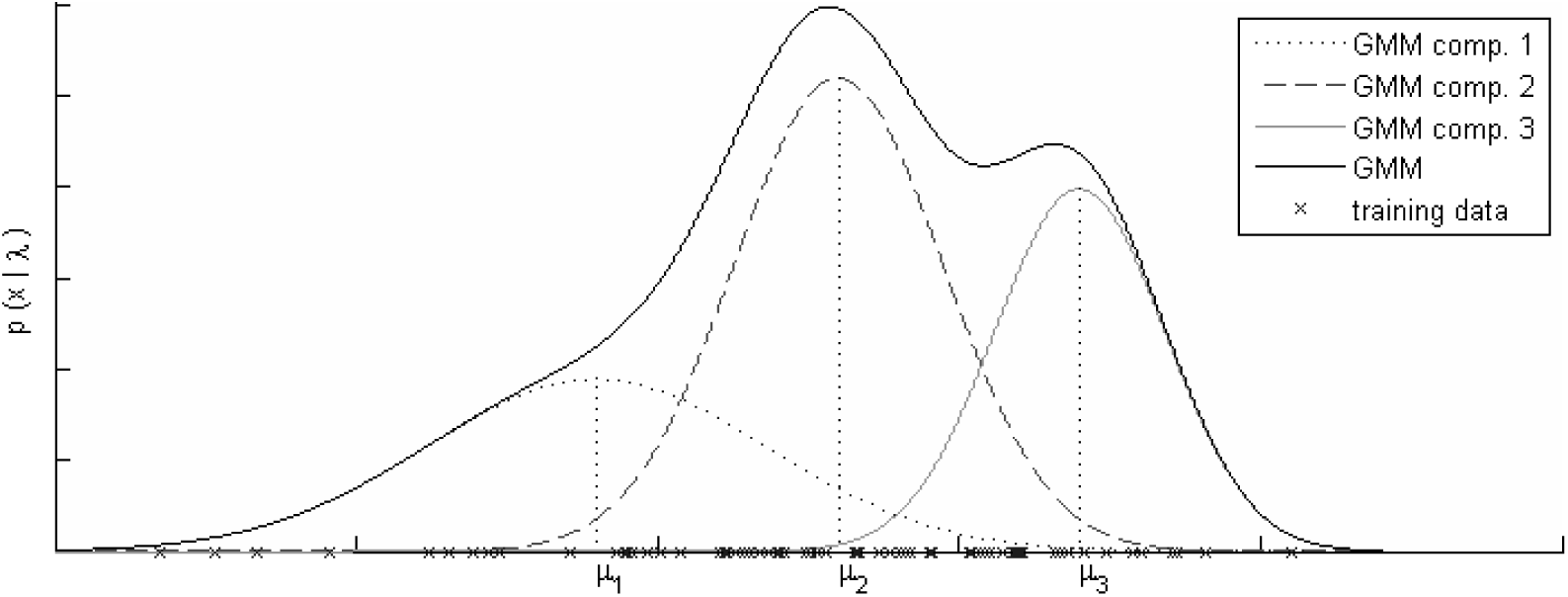
Given a set of one dimensional feature vectors (x-axis), the Gaussian Mixture Model with three mixture components which best describes the data set (in the sense of maximal likelihood (4)) is given by the solid line. Note that the GMM is formed from 3 normal distributions each weighted by the relative number of vectors it encloses (Machlica 2012).

#### Mashona mole-rat

The Mashona mole-rat (*Fukomys darlingi*), formerly known as *Cryptomys*, belongs to African endemic rodent family Bathyergidae see Figure 4. This herbivorous species inhabits Miombo woodland and shrub habitats in eastern and northern Zimbabwe, northern Mozambique and southern Malawi (Bennett and Faulkes 2000). Mashona mole-rat lives in underground burrows in small families of 5-9 individuals (Bennett et al. 1994). The families are composed of a reproductive pair and a number of non-reproductive descendants (Bennett et al. 1994). The colony works under strict hierarchy in which the reproductive individuals are usually the most dominant over others of the respective gender (Gebathuler et al. 1996).

**Figure 4:**
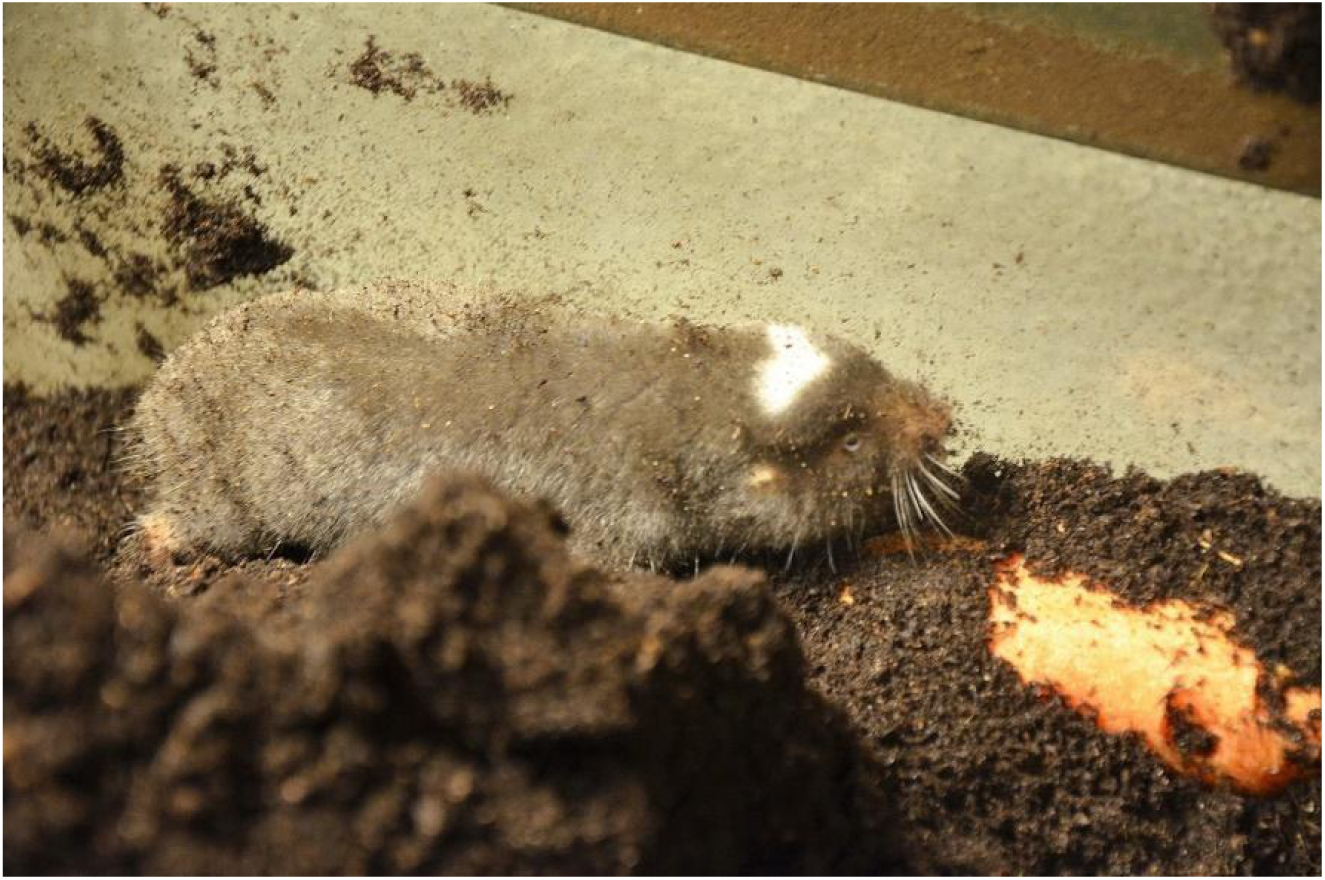
The Mashona mole-rat individual.

Mashona mole-rats are strictly subterranean rodents who mate, breed and forage underground and rarely come to surface (Begall et al. 2007). Living in underground ecotope brings special requirements on sensory capabilities of its inhabitants and favors some types of communication over others (reviewed in Burda et al. 1990, Francescoli 2000). Absence of light in burrows prevents visual communication. Mole-rats eyes are reduced and degenerated (Burda et al. 1990, Cernuda-Cernuda et al. 2003, Nemec et al. 2004, 2008, Hetling et al. 2005) and can be used to perceive light, to maintain circadian activity and to avoid burrow breaches (Němec et al. 2007; Kott et al. 2010). Poor vision can be partly compensated by tactile sense that is well developed in subterranean rodents. However, it is only useful to close proximity of stimuli (Burda *et al*. 1990, Park *et al*. 2007). Furthermore reduced airflow in burrows negatively affects olfactory sense because the dispersal of scent signals is limited. On the other hand acoustic signals can propagate over distances of several meters within natural burrows (Hethet al. 1986; Lange et al. 2007). Under such conditions, vocalization becomes a crucial means of communication and therefore widely studied in subterranean rodents. In case of Mashona mole-rats 12 types of acoustic signals consisting of 10 true vocalizations and 2 mechanically produced sounds were described (Dvořáková et al. 2016).

#### Vocalization

This species possess two types of mating calls; see Figure 5, both emitted mostly by females during courtship (Dvořáková et al. 2016). These calls are often produced in a series when one type alternates the other. A cluck is a very short vocalization, with the mean duration of 0.03 s. The range of frequency is very low and it usually does not exceed 5 kHz. A shriek is a sound similar to a cluck, but it has a main frequency lower than a cluck and does not show a rising frequency towards the end.

**Figure 5:**
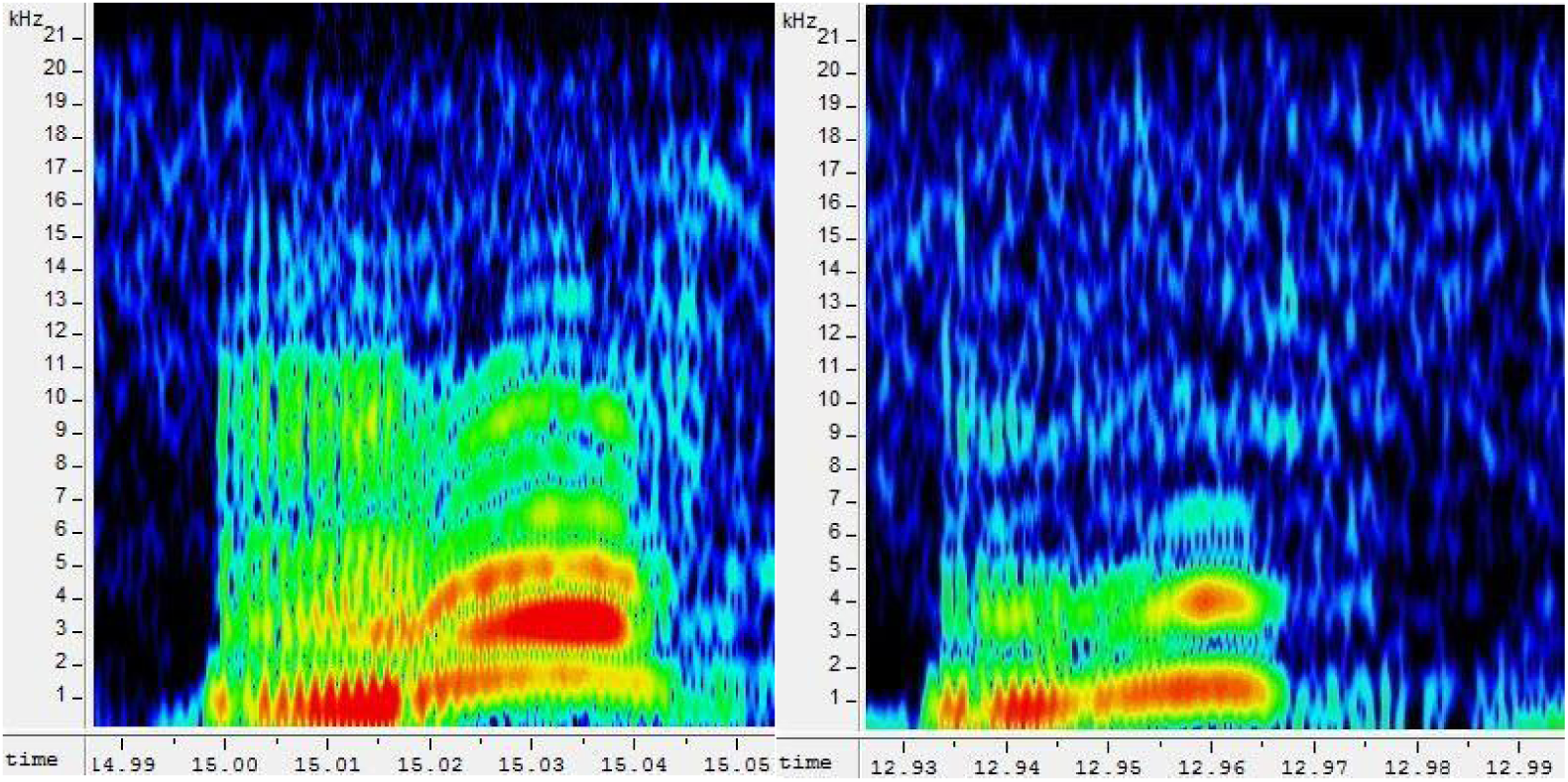
Spectrograms of the mating calls: cluck (left), and shriek (right).

#### Data

From the vocal repertoire of Mashona mole-rats mating calls were selected as the type of vocalization to most likely carry informations about the signallers identity. The recordings were taken with the MD 431 II Sennheiser dynamic microphone (frequency range 40-16.000 Hz) and recorded with the Marantz card audio recorder PMD660 (sample frequency 44.1 kHz, resolution 16 bit). The mole-rats were simultaneously recorded using a Panasonic SDRH60EP-S camera to enable repeated analysis of the testing sessions.

For individual identification, vocalizations of five dominant (breeding) females were used. The recordings were divided into 20-second tracks. Each soundtrack was named by a combination of a single letter (A,B,..E) represents a particular female, and a number represents a no. oftrack, e.g. “A_04”, or “E_11”.

Families or pairs were kept in terrariums with horticultural peat used as substrate and supplemented with plastic tubes as imitations of tunnels and flowerpots to simulate the nest see Figure 6. University of South Bohemia, Faculty of Science has mole-rat breeding facility that houses one of the most representative collection of the underground mammals in the world. The experiments were carried out on these captive animals and they serve to verify and add wild nature findings. The room was lighted in 12D/12L (lights on at 0700 h). The temperature was kept at 25±1 °C. Animals were fed *ad libitum* with carrots, potatoes, apples and dry rodent food.

**Figure 6:**
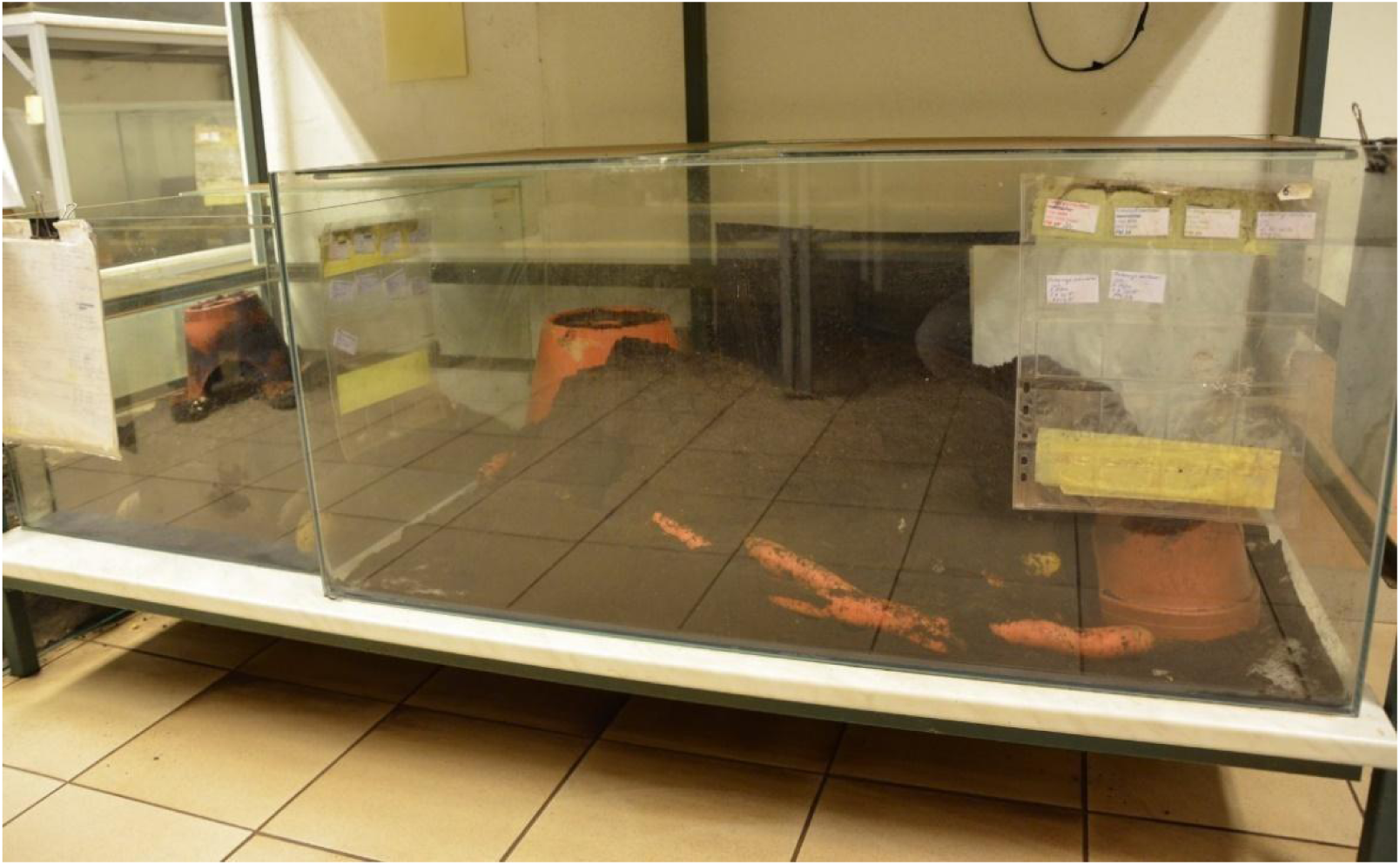
The Mashona mole-rat breeding facility in the University of South Bohemia, Faculty of Science.

#### Experiment evaluation, Equal Error Rate

Four different situations may occur during the verification, see Figure 7.

**Figure 7:**
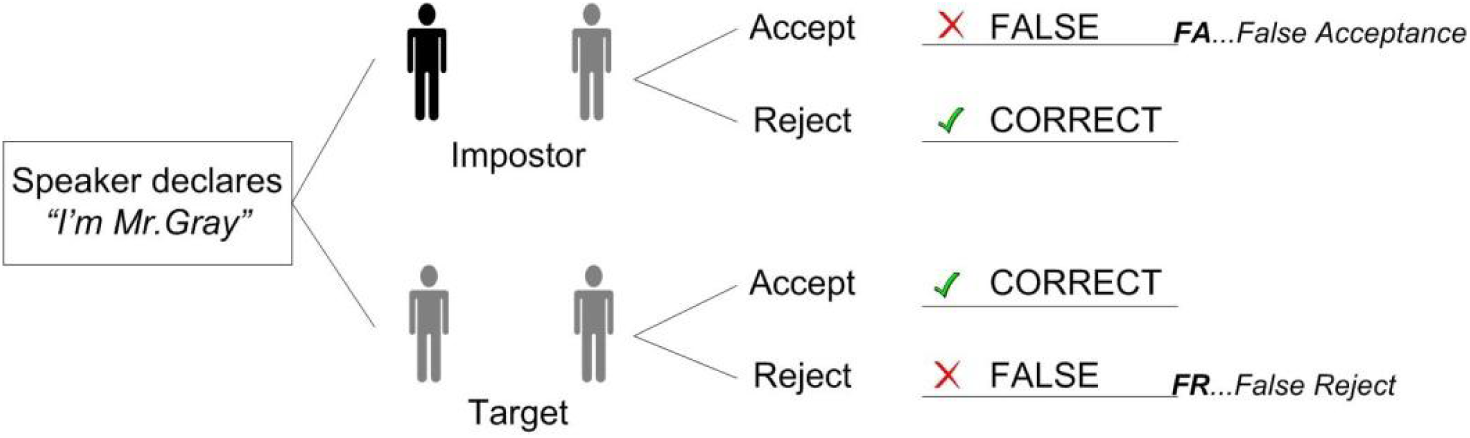
Verification, false and correct decision.

Incorrect acceptance error *R*_*FA*_(*Θ*) is defined as

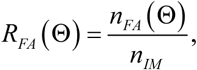

where *Θ* is the threshold (see below), *n*_*FA*_ is the number of cases when the system incorrectly accepts the impostor, and *n*_*IM*_ is the total number of cases where an impostor has been tested.

Incorrect rejection error *R*_*FR*_(*Θ*) is defined as

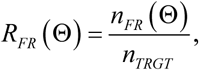

where *n*_*FR*_ is the number of cases when the system incorrectly rejected the Target (right speaker/bird) and *n*_*TRGT*_ is the total number of cases where the target has been tested. Setting the threshold *Θ* affects the total number of *R*_*FA*_ and *R*_*FR*_. Increasing the threshold reduces the false acceptance error rate *FA*, but it simultaneously increases the false rejection *FR* error. This happens because the system requires a higher probability of similarity. On the contrary, if the threshold is lower, the *FR* error decreases, but the *FA* increases as the system needs lower probability of similarity to accept the speaker. This leverage effect is summarized in Table 1. Both errors are called *operating point* (Bimbot 2004).

**Table 1:** Level of threshold *Θ* value and error rates.

|  |  |
| --- | --- |
| Highest $\Theta$ | $R_{FA}(\Theta)$ decrease |
| | $R_{FR}(\Theta)$ increase |
| Lowest $\Theta$ | $R_{FA}(\Theta)$ increase |
| | $R_{FR}(\Theta)$ decrease |

The *Equal Error Rate* (*EER*) is used for single number evaluation of the system, which indicates the threshold value *Θ*_*EER*_ at which *R*_*FA*_ and *R*_*FR*_ are equal. It is defined as

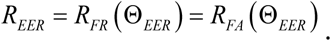

In real experiments, however, a threshold *Θ* must be set first, where after the decisions the *R*_*FA*_ and *R*_*FR*_ errors can be calculated. Finding the threshold *Θ*_*EER*_ can therefore be nontrivial.

### Experiments

#### Testing procedure

As stated in section Methods, the experiments were performed by the framework based on the GMM-UBM method, described by (Ptáček 2016). For individual recognition, vocalizations of five dominant (breeding) females were selected. The recordings were divided into 20-second soundtracks.

During the verification process the framework performs *trials*. In each *trial* two soundtracks are compared: the trained one (which is included in the GMM set) and tested one (which is included in the TEST set). Framework has to decide whether the TEST soundtrack do belong or do not belong to the trained (GMM) animal. The verification system works on basis of Yes or No decisions. Since the TEST animal could be the the trained one (GMM) or not (so called *impostor*), two types of errors could occur, see Figure 7.

- False acceptance…the trained and tested individuals are no the same, but the framework wrongly evaluated the animals as the same.
- False rejection…, the trained and tested individuals are the same, but the framework wrongly evaluated the animals as not the same.

The success rate gives a ratio of errors and correct answers. To ensure the objectivity of testing, two rounds were performed each female where the UBM set was filled differently.

We also included four older recordings from female B (recorded in 2010). These four recordings were of poor quality as they were recorded by a different type of audio recorder (Sony Digital Audio Tape-corder TCD-D100) and probably negatively affected the results when used for GMM model estimating of this female B. On the other hand, these recordings did not have any impact when used for UBM sets and in the verification phase. To give reasonable results we applied these recordings just for the TEST and UBM sets, not for GMM model training.

#### Parameters set up

At first, the suitable parameters were analysed because there is not any recommended parameters set up for an animal vocalization song unlike for human speech (for instance, window length 30 ms, overleap 15 ms, pre-emphasis 0.97, etc.). Actually, the setup has to be evaluated from the scratch for each research.

The Framework cannot perform a multi-dimensional optimization. Therefore, this process is complicated because there are many crucial parameters, which have to be tuned one at a time. See iteration steps in Table 2 as an example of parameters optimization process.

**Table 2:** Example of the parameters optimization approach.

| Iteration | Windows |  | Filters |  | Frequency |  | VAD | Threshold | Accuracy |
| --- | --- | --- | --- | --- | --- | --- | --- | --- | --- |
| # | Length | Shift | Mel On/Off | Number | Low | High | Automatic sound segmentation | - | Success rate of identification |
| 1 | 30 | 20 | off | 5 | 2500 | 20000 | off | 5,2 | 75% |
| 2 | 30 | 20 | off | 5 | 200 | 10000 | off | 3,3 | 49% |
| 3 | 30 | 20 | off | 5 | 100 | 20000 | off | 4,5 | 61% |
| 4 | 20 | 10 | off | 5 | 100 | 20000 | off | 4,1 | 56% |
| 5 | 20 | 10 | off | 5 | 2000 | 20000 | off | 3,9 | 71% |
| 6 | 20 | 10 | off | 5 | 1000 | 20000 | off | 3,65 | 60% |
| 7 | 40 | 20 | off | 5 | 1000 | 20000 | off | 4,4 | 63% |
| 8 | 40 | 20 | off | 5 | 2000 | 10000 | off | 2,67 | 65% |
| 9 | 20 | 10 | off | 25 | 500 | 19000 | off | 0,19 | 62% |
| 10 | 20 | 10 | off | 25 | 500 | 19000 | on | 0,09 | 65% |
| 11 | 20 | 10 | off | 25 | 500 | 19000 | on | 1,38 | 64% |
| 12 | 20 | 10 | on | 25 | 500 | 19000 | on | 0,28 | 79% |
| 13 | 20 | 10 | off | 25 | 100 | 19000 | on | 0,45 | 66% |
| 14 | 20 | 10 | off | 25 | 500 | 20000 | on | 0,36 | 76% |
| 15 | 20 | 10 | on | 25 | 100 | 20000 | on | 0,24 | 75% |
| 16 | 20 | 10 | on | 25 | 250 | 20000 | on | 0,282 | 80% |
| 17 | 20 | 10 | on | 25 | 50 | 20000 | on | 0,19 | 77% |
| 18 | 30 | 15 | on | 25 | 500 | 20000 | on | 1,64 | 78% |

Final values of the main parameters after optimization are cited in Table 3.

**Table 3:** Parametrization set up values.

| # | Parameter | Value |
| --- | --- | --- |
| 1 | Window type | Hamming |
| 2 | Window length | 20 ms |
| 3 | Window overlap | 10 ms |
| 4 | Number of filters | 23 |
| 5 | Number of cepstral coefficients | 20 |
| 6 | Compute zero coefficient logE | Yes |
| 7 | High pass filter | 250 Hz |
| 8 | Low pass filter | not used |
| 9 | Delta coefficients | Yes |
| 10 | VAD detector | No |
| 11 | Preemphasis | 0.99 |
| 12 | Linear/MelFilter scale | Mel |

#### Results

Table 4 shows the overall success rate of individual identification. The number of recordings (20 s soundtracks) varied from 10 to 40.

**Table 4:**
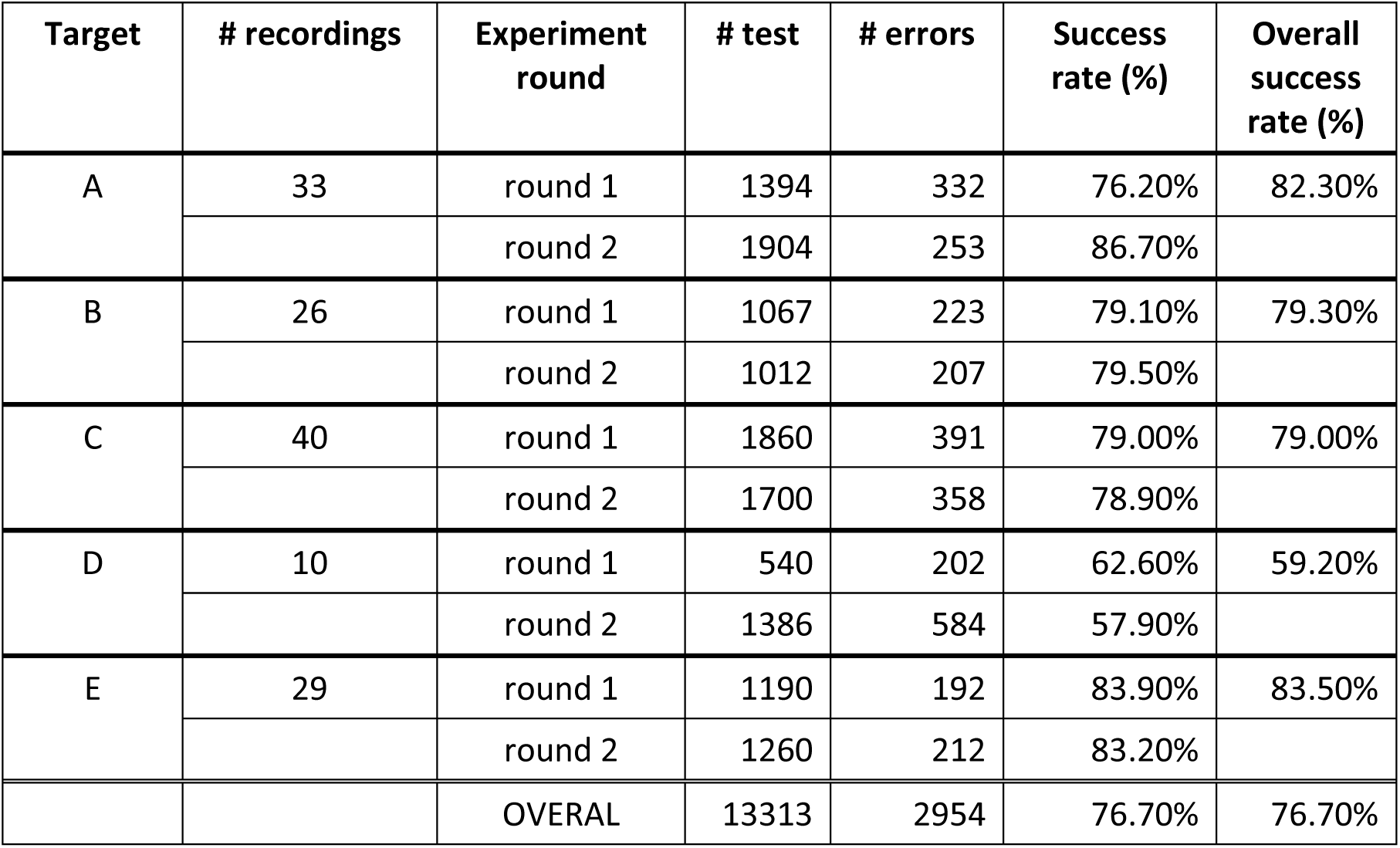
Mole-rat identification results.

| Target | # recordings | Experiment round | # test | # errors | Success rate (%) | Overall success rate (%) |
| --- | --- | --- | --- | --- | --- | --- |
| A | 33 | round 1 | 1394 | 332 | 76.20% | 82.30% |
|  |  | round 2 | 1904 | 253 | 86.70% |  |
| B | 26 | round 1 | 1067 | 223 | 79.10% | 79.30% |
|  |  | round 2 | 1012 | 207 | 79.50% |  |
| C | 40 | round 1 | 1860 | 391 | 79.00% | 79.00% |
|  |  | round 2 | 1700 | 358 | 78.90% |  |
| D | 10 | round 1 | 540 | 202 | 62.60% | 59.20% |
|  |  | round 2 | 1386 | 584 | 57.90% |  |
| E | 29 | round 1 | 1190 | 192 | 83.90% | 83.50% |
|  |  | round 2 | 1260 | 212 | 83.20% |  |
|  |  | OVERAL | 13313 | 2954 | 76.70% | 76.70% |

The success rate of correct identification of individual varied between 59.2% (female D) and 83.5% (female E). For particular experiments, the success rate varied between 57.9% (female D) and 86.7% (female A). The lowest number of correct identification was obtained for female D, which unfortunately died at the time of recordings. We obtained only 10 soundtracks from this female, which is significantly less compared to any other female (see Table 4, second row).

Table 5 defines the assembly of the three data sets (GMM, UBM, TEST) for each experiment round. To quote this distribution is needed for experiment repetition if required. Notice, in case animal is included in both the GMM and TEST sets, the recordings are different as stated in section Experiment evaluation.

**Table 5:**
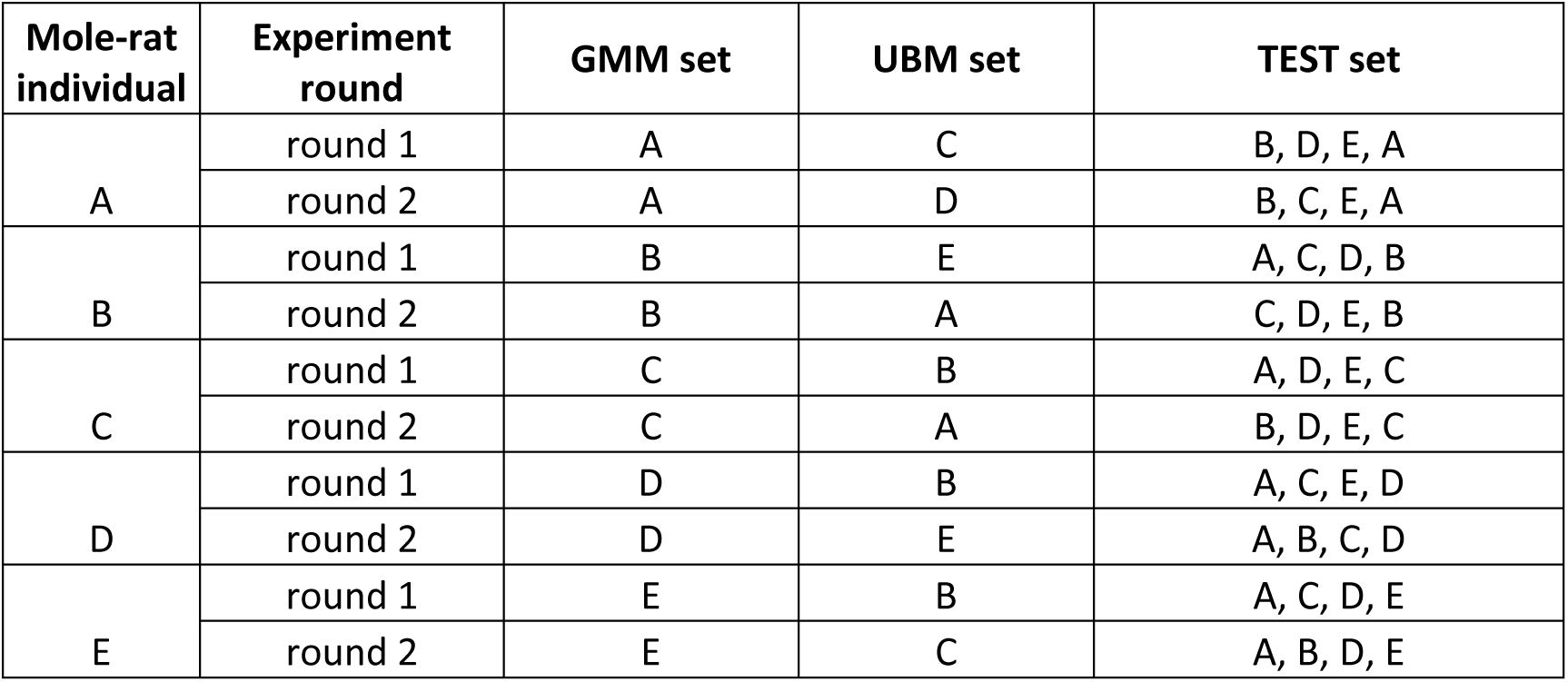
Assembly of the three data sets (GMM, UBM, TEST). Notice, in case animal is included in both the GMM and TEST sets, the recordings are different.

## Conclusion

In this study the programmed framework automatically verified the mole-rat individuals on its vocalization. The aim of these experiments was to test a mole-rat vocalization for presence of diverse individually distinctive features. The GMM-UBM automatic framework achieved identification accuracy 76.7%, the lowest 59.2%, and the highest 83.5%.

The experiments confirmed the hypothesis that the mole-rats’ vocalization also holds individuality identification (Bednářová et al. 2013). As described above, this hypothesis is of high importance because of the mole-rats’ life environment. Based at our knowledge, these are the first experiments dealing with mole-rat automatic individual identification.

The overall percentage proves that the mating calls of the Mashona mole-rat can carry information about mole-rat individuality. Our results showed that studied vocal signals in the Mashona mole-rats are individually specific which indicates the possibility of individual vocal recognition in this species (Dvořáková 2013).

## Acknowledgement

This publication was supported by the project LO1506 of the Czech Ministry of Education, Youth and Sports.

